# DEET and other repellents are inhibitors of mosquito odorant receptors for oviposition attractants

**DOI:** 10.1101/711481

**Authors:** Pingxi Xu, Fangfang Zeng, Robert H. Bedoukian, Walter S. Leal

## Abstract

In addition to its primary function as an insect repellent, DEET has many “off-label” properties, including a deterrent effect on attraction of gravid female mosquitoes. DEET negatively affects oviposition sites. While deorphanizing odorant receptors (ORs) using the *Xenopus* oocyte recording system, we have previously observed that DEET generated outward (inhibitory) currents on ORs sensitive to oviposition attractants. Here, we systematically investigated these inhibitory currents. We recorded dose-dependent outward currents elicited by DEET and other repellents on ORs from *Culex quinquefasciatus, Aedes aegypti*, and *Anopheles gambiae*. Similar responses were observed with other plant-derived and plant-inspired compounds, including methyl jasmonate and methyl dihydrojasmolate. Inward (regular) currents elicited by skatole upon activation of CquiOR21 were modulated when this oviposition attractant was coapplied with a repellent. Compounds that generate outward currents in ORs sensitive to oviposition attractants elicited inward currents in a DEET-sensitive receptor, CquiOR136. The best ligand for this receptor, methyl dihydrojasmolate, showed repellency activity, but was not as strong as DEET in our test protocol.

## 1. Introduction

Insect repellents have been used since antiquity as prophylactic agents against diseases transmitted by mosquitoes and other arthropods. They developed gradually from smoke generated by burning plants (e.g., lemon gum) and topical applications of essential oils (e.g., lemon eucalyptus extract) into repellent substances, including those isolated from plants (e.g., p-menthane-3,8-diol, PMD) and a broad-spectrum synthetic repellent DEET (*N,N*-dimethyl-3-methylbenzamide), which was discovered in the early 1940s from a screening of more than 7000 compounds (Moore and Debboun, 2006). Thereafter, other synthetic repellents have been developed, including IR3535, (ethyl-3-(N-n-butyl-N-acetyl)aminopropionate) and picaridin (butan-2-yl 2-(2-hydroxyethyl)piperidine-1-carboxylate) (Boeckh et al., 1996), but DEET remains the most widely used repellent substance worldwide (Moore and Debboun, 2006), particularly in the United States of America.

Repellents work primarily as spatial and contact repellents. Mosquitoes attracted to and flying towards vertebrate hosts (e.g., humans) may make oriented movements away from the source upon approaching chemically treated skin surfaces. In this case, the chemical is a repellent sensu stricto (Dethier et al., 1960). Because the repellent is acting from a distance (in the vapor phase [Barton-Browne, 1977]), it may be referred to as a spatial repellent (Gouck et al., 1967). When mosquitoes land on a chemically treated skin thus making contact before starting increasing locomotion activity or taking off, the chemical is called a contact repellent, which is sometimes referred to as excitorepellent, irritant, or contact irritant (Grieco et al., 2007). From a strict mechanistic viewpoint, these two groups of compounds should be named noncontact and contact disengagent (Miller et al., 2009) for spatial and contact repellents, respectively. It is now known that at least *Culex* and *Aedes* mosquitoes smell DEET (Stanczyk et al., 2010; Syed and Leal, 2008). More importantly, it has been demonstrated that an odorant receptor (OR) from the Southern house mosquito, *Cx. quinquefasciatus*, CquiOR136, is essential for reception of DEET as a noncontact disengagent (Xu et al., 2014). Recently, it has been demonstrated that as a contact disengagent, DEET is detected by sensilla on the tarsal segments of the legs of the yellow fever mosquito *Aedes (=Stegomya) aegypti* (Dennis et al., 2019), but the receptors remain elusive. Lastly, it has been suggested that DEET merely masks the reception of human emanations by *Anopheles coluzzii (=An. gambiae M form)* (Afify et al., 2019), thus reducing the attractiveness of the host.

Although its modes of action remain a matter of considerable debate, DEET is a gold-standard repellent. It also has many “off-label” properties that do not directly affect human-mosquito interactions. For example, DEET is a feeding deterrent (Lu et al., 2017), but if this were the primary mode of action it would have little value in epidemiology. The great value of repellents is that they reduce biting rates, which represents a second order parameter in vector capacity (Norris and Coats, 2017). Another property that may have a value in epidemiology, albeit not by decreasing vector capacity, is the deterrent effect of DEET on oviposition, as first observed for *Ae. aegypti* (Kuthiala et al., 1992).

While using the *Xenopus* oocyte recording system to deorphanize odorant receptors (ORs) involved in the reception of oviposition attractants, we observed that DEET elicited outward currents in our preparations, in contrast to oviposition attractants and other compounds that generated inward (regular) currents. We have now systematically investigated this phenomenon using different ORs from three different species of mosquitoes. Here, we report that DEET, IR3535, and picaridin elicit outward (inhibitory) currents on OR involved in the reception of mosquito oviposition attractants in the Southern house mosquito, *Cx. Quinquefasciatus*, and orthologues from the yellow fever and malaria mosquitoes. Dose-dependent outward currents were also observed with compounds in a panel that included plant-derived and plant-inspired repellents. Like DEET, IR3535 and picaridin (Xu et al., 2014), plant-inspired compounds, elicited robust inward current in the DEET receptor, CquiOR136, and showed repellency activity.

## 2. Materials and Methods

### 2.1. AaegOrco cloning

The pGEM-HE plasmids for the following ORs were obtained as previously reported: CquiOrco (Hughes et al., 2010), CquiOR21 (Pelletier et al., 2010), CquiOR2 (Hughes et al., 2010), CquiOR37, and CquiOR99 (Zhu et al., 2013). pSP64 Poly (A) or pT7TS vectors carrying AgamOrco (Pitts et al., 2004), AgamOR10 (Carey et al., 2010; Wang et al., 2010), AgamOR8 (Lu et al., 2007), AgamOR40 (Liu et al., 2010), and AaegOR10 (Bohbot et al., 2007) were generously shared by Dr. Larry Zwiebel, Vanderbilt University. To obtain a full-length coding sequence of AaegOrco, total RNA was extracted from *Ae. aegypti* female mosquitos provided by Dr. Anthon J. Cornel, UC Davis, Department of Entomology and Nematology, by using TRIzol (Invitrogen, Carlsbad, CA). cDNA was synthetized from 1 μg of total RNA using a GoScript™ Reverse Transcript kit, according to the manufacturer’s manual (Promega, Madison, WI). Then, we performed PCR using AaegOrco gene-specific primers, AaegOrco-F 5’-accATGAACGTCCAACCGACAAAGTACCATG-3’ with a Kozak sequence, AaegOrco-R 5’-TTATTTCAACTGCACCAACACCATGAAGTAGG-3’. The gene was cloned into pGEM-HE vector through the In-Fusion HD Cloning system (Clontech, Mountain View, CA). Amino acid sequence was identical to that in VectorBase.

### 2.2. In Vitro Transcription, Oocytes Microinjection, and Electrophysiology

In vitro transcription, oocytes microinjection, and electrophysiology were performed as previously described (Xu et al., 2014). Briefly, in vitro transcription of cRNAs was performed by using an mMESSAGE mMACHINE T7 kit (Ambion), according to the manufacturer’s protocol. Plasmids were linearized with NheI, SphI, or PstI, and capped cRNAs were transcribed using T7 or SP6 RNA polymerase. cRNA samples were purified with LiCl precipitation solution and resuspended in nuclease-free water at a concentration of 200 μg/mL and stored at −80°C in aliquots. RNA concentrations were determined by UV spectrophotometry. cRNA samples were microinjected into stage V or VI *Xenopus laevis* oocytes (EcoCyte Bioscience, Austin, TX). Oocytes were then incubated at 18°C for 3–7 days in modified Barth’s solution [in mM: 88 NaCl, 1 KCl, 2.4 NaHCO_3_, 0.82 MgSO_4_, 0.33 Ca(NO_3_)_2_, 0.41 CaCl_2_, 10 HEPES, pH 7.4] supplemented with 10 μg/mL of gentamycin, 10 μg/mL of streptomycin, and 1.8 mM sodium pyruvate. A two-electrode voltage clamp (TEVC) was used to detect currents. Oocytes were placed in a perfusion chamber (flow rate was 10 mL/min) and challenged with test compounds. Odorant-induced currents were amplified with an OC-725C amplifier (Warner Instruments, Hamden, CT), voltage held at −80 mV, low-pass filtered at 50 Hz and digitized at 1 kHz. Data acquisition and analysis were carried out with Digidata 1440A and pClamp10 software (Molecular Devices, LLC, Sunnyvale, CA).

### 2.3. Panel of Odorants

The following compounds were tested: skatole (CAS# 83-34-1), fenchone (CAS# 1195-79–5), 1-octen-3-ol (CAS# 3391-86-4), DEET (CAS# 134-62-3), IR3535 (CAS# 52304-36-6), PMD (CAS# 42822-86-6), picaridin (CAS# 119515-38-7), BDR-1 (farnesyl cyclopentanone, CAS# not available, n/a), BDR-2 ((*E,E*)-farnesol, CAS# 106-28-5), BDR-3 (methyl dihydrojasmonate = hedione, CAS# 24851-98-7), BDR-4 (methyl jasmonate, CAS# 39924-52-2), BDR-5 (γ-dodecalactone, CAS# 2305-05-7), BDR-6 (δ-tetradecalactone, CAS# 2721-22-4), BDR-7 (ethyl palmitate, CAS# 628-97-7), BDR-8 (isophorol, CAS# 470-99-5), BDR-9 (isophorone, CAS# 78-59-1), BDR-10 (prenyl dihydrojasmonate, CAS# n/a), BDR-11 (2-pentadecanol, CAS# 1653-34-5), BDR-12 (3,5,5-trimethyl cyclohexanol, CAS# 116-02-9), BDR-13 (methyl apritol, CAS# n/a), BDR-14 (methyl dihydrojasmolate, CAS# n/a), BDR-15 (dihydrojasmonic acid, CAS# 3572-64-3), BDR-16 (methyl apritone = miranone, 1206769-45-0), BDR-17 (dihydrojasminlactone, CAS# n/a), BDR-18 (dihydrojasmindiol, CAS# n/a), BDR-19 (ethyl dihydrojasmonate, CAS# n/a), and BDR-20 (2-pentadecanone, CAS#2345-28-0). To avoid possible mislabeling, after sample preparation for electrophysiology and behavior identity of test chemicals was confirmed by gas chromatography-mass spectrometry (GC-MS) using a 5973 Network Mass Selective Detector linked to a 6890 GC Series Plus + (Agilent Technology, Palo Alto, CA). The GC was equipped with an HP-5MS capillary column (30 m × 0.25 mm; 0.25 μm, Agilent Technologies), which was operated at 70°C for 1 min and increased at a rate of 10°C/min to 270°C, with a final hold of 10 min and a post run of 10 min at 290°C.

### 2.4. Mosquito Repellency Assay

The surface landing and feeding assay has been detailed elsewhere (Leal et al., 2017; Xu et al., 2014). In short, two 50-mL Dudley bubbling tubes painted internally with black hobby and craft enamel (Krylon, SCB-028) were held in a wooden board (30 x 30 cm), 17 cm apart from each end and 15 cm from the bottom. The board was attached to the frame of an aluminum collapsible field cage (30.5 × 30.5 × 30.5 cm; Bioquip, Rancho Cordova, CA, USA). Two small openings were made 1 cm above each Dudley tube to hold two syringe needles (Sigma-Aldrich, 16-gauge, Z108782) to deliver CO_2_. To minimize handling of mosquitoes, test females had been kept inside collapsible field cages since latest pupal stage. These female cages had their cover premodified for behavioral studies. A red cardstock (The Country Porch, Coeur d’Alene, ID, GX-CF-1) was placed internally at one face of the cage, openings were made in the cardboard and in the cage cover so the cage could be attached to the wooden board with the two Dudley tubes and CO_2_ needles projecting inside the mosquito cage 6 and 3 cm, respectively. Additionally, windows were made on the top and the opposite end of the red cardstock for manipulations during the assays and a video camera connection, respectively. The two cages were connected at least 2 h prior to bioassays. At least 10 min before the assays, water at 28°C started to be circulated with a Lauda’s Ecoline water bath, and CO_2_ at 50 mL/min was delivered from a gas tank just at the time of the behavioral observations. Sample rings were prepared from strips of filter papers 25 cm-long and 4-cm wide and hung on the cardstock wall by insect pins to make a circle around the Dudley tubes. Cotton rolls (iDental, Fort Worth, TX, 1 x 3 cm) were loaded with 100 μl of defibrillated sheep blood purchased from UC Davis VetMed shop and placed between a Dudley tube and CO_2_ needle. For each run, one paper ring was loaded with 200 μL of hexane (control) and the other with 200 μL of test repellent (DEET or methyl dihydrojasmolate) in hexane. Solvent was evaporated for 1-2 min, blood-impregnated cotton plugs and filter paper rings were placed on the arena, CO_2_ was started, and the assays recorded with an infrared camera (Sony Digital Handycan, DCR-DVD 810). During the assay, the arena was inspected with a flashlight with red filter. After 5 min, the number of females that have landed and continued to feed on each side of the arena was recorded. Insects were gently removed from the cotton rolls and the assays re-initiated after rotation of sample and control. Thus, repellence for each set of test mosquitoes was measured with the filter paper impregnated with the same sample at least once on the left and once on the right side of the arena.

### 2.5. Graphic Preparations and Statistical Analysis

Graphic illustrations were prepared with Prism 8 (GraphPad, La Jolla, CA). Number of mosquitoes in the treatment (T) and control (C) side of the arena were transformed into *%* protection, P% = (1-[T/C]) x 100, according to WHO (WHO, 2009) and EPA (EPA, 2010) recommendations. Tests comparing two repellents were conducted in tandem, with two replicates for DEET (treatment right and then left or left and then right), followed by two replicated from a test repellent (BDR-14), with this cycle being repeated multiple times. Data that passed the Shapiro-Wilk normality test were analyzed with the two-tailed, unpaired *t* test; otherwise, data were analyzed with the Mann-Whitney test. All data are expressed as mean ± SEM.

## 3. Results and Discussion

### 3.1 Repellent-elicited outward currents

To revisit our earlier observation of repellent-induced outward currents on OR sensitive to oviposition attractants, we challenged CquiOR21/CquiOrco-expressing oocytes with DEET and then skatole. CquiOR21, formerly known as CquiOR10 (Leal et al., 2013), is narrowly tuned to the oviposition attractant skatole (Hughes et al., 2010). CquiOR21/CquiOrco-expressing oocytes generated robust inward (regular) currents when stimulated with 10 μM skatole, whereas 1 mM DEET elicited outward currents (Fig. 1). These outward currents were dose-dependent (Fig. S1A) and were not observed when oocytes were injected only with CquiOrco cRNA (Fig. S1B) or CquiOR21 cRNA (Fig. S1C).

**Fig. 1.**
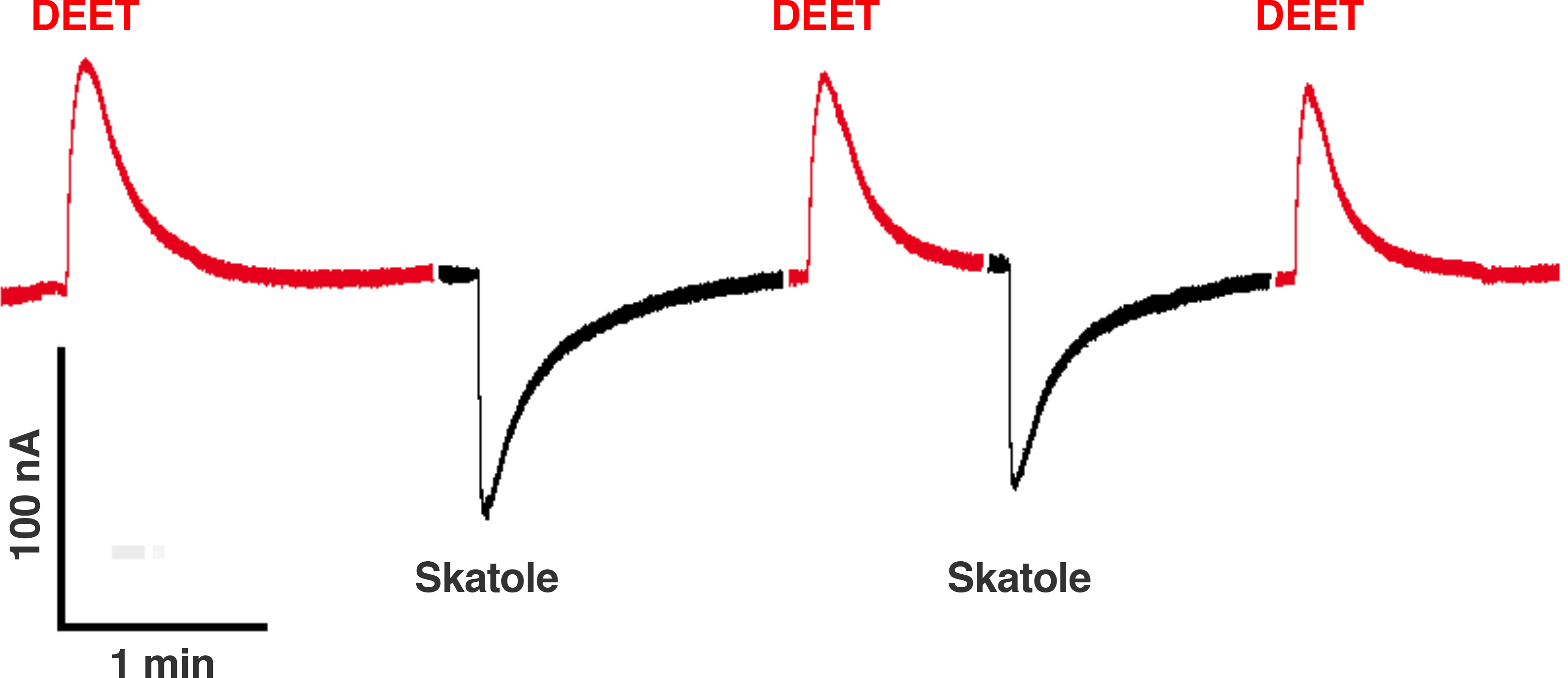
Responses of a CquiOR21·CquiOrco-expressing oocyte to DEET and skatole. Outward and inward currents elicited by 1 mM DEET (red traces) and 0.1 μM skatole (black traces), respectively.

We then tested how CquiOR21 would respond to other commercially available repellents, i.e., PMD, IR3535, and picaridin. In these new preparations, CquiOR21/CquiOrco-expressing oocytes responded to DEET and IR3535 with dose-dependent outward currents (Fig. 2A). Picaridin elicited minor outward currents at lower doses but robust outward currents at 1 mM dose. By contrast, PMD did not elicit outward currents; it was silent at lower doses and gave minor inward currents at the highest dose, 1 mM (Fig. 2A). We then interrogated CquiOR21 orthologs from the yellow fever mosquito, AaegOR10 (Bohbot et al., 2007), and the malaria mosquito, AgamOR10 (Carey et al., 2010; Wang et al., 2010). AaegOR10/AaegOrco- and AgamOR10/AgamOrco-expressing oocytes responded with a similar pattern to that observed with CquiOR21/CquiOrco-expressing oocytes (Fig. 2B, C). Specifically, DEET generated dose-dependent outward currents as did picaridin at 1 mM, whereas PMD elicited only minor currents. Over the years, we have deorphanized multiple ORs from *Cx. quinquefasciatus* and were surprised to observe that these outward currents generated only with preparations involving ORs sensitive to oviposition attractants. We then tested other ORs for oviposition attractants, namely CquiOR121 (=CquiOR2) (Leal et al., 2013; Pelletier et al., 2010), CquiOR37, and CquiOR99 (Zhu et al., 2013). Oocytes expressing each of these ORs along with the obligatory coreceptor Orco elicited dose-dependent outward currents when challenged with DEET (Fig. S2). We challenged also other ORs from the malaria mosquito, which are not involved in the reception of oviposition attractants. Like their *Culex* counterparts, ORs unrelated to oviposition attractants did not generate outward currents when challenged with DEET (Fig. S3). A previously reported larval OR, AgamOR40 (Liu et al., 2010) generated dose-dependent inward currents in response to DEET as well as to its best ligand, fenchone (Fig. S3A). By contrast, oocytes expressing AgamOR8 (Bohbot and Dickens, 2009; Lu et al., 2007) along with AgamOrco generated robust, dose-dependent inward currents in response to 1-octen-3-ol, but it was silent to DEET (Fig. S3B). Previously, it has been demonstrated that DEET modulates responses of other odorants to ORs (Bohbot and Dickens, 2010), but no outward currents were recorded. When odorants were present in combination with DEET at high doses, the odorant-induced inward currents decreased significantly (Bohbot and Dickens, 2010), but DEET per se did not elicite measurable currents. At the time of this writing, a small DEET-induced hyperpolarization of a mosquito OR was reported (Dekel et al., 2019). Our fortuitous discovery of outward current elicited by DEET might occur mainly on ORs sensitive to oviposition attractants. However, we have recently reported outward currents elicited by multiple compounds, including repellents, on a *Culex* OR, CquiOR32, which is sensitive to a plant-derived compound with repellency activity, methyl salicylate (Xu et al.). It is, therefore, conceivable that the phenomenon expands beyond OR sensitive to mosquito oviposition attractants.

**Fig. 2.**
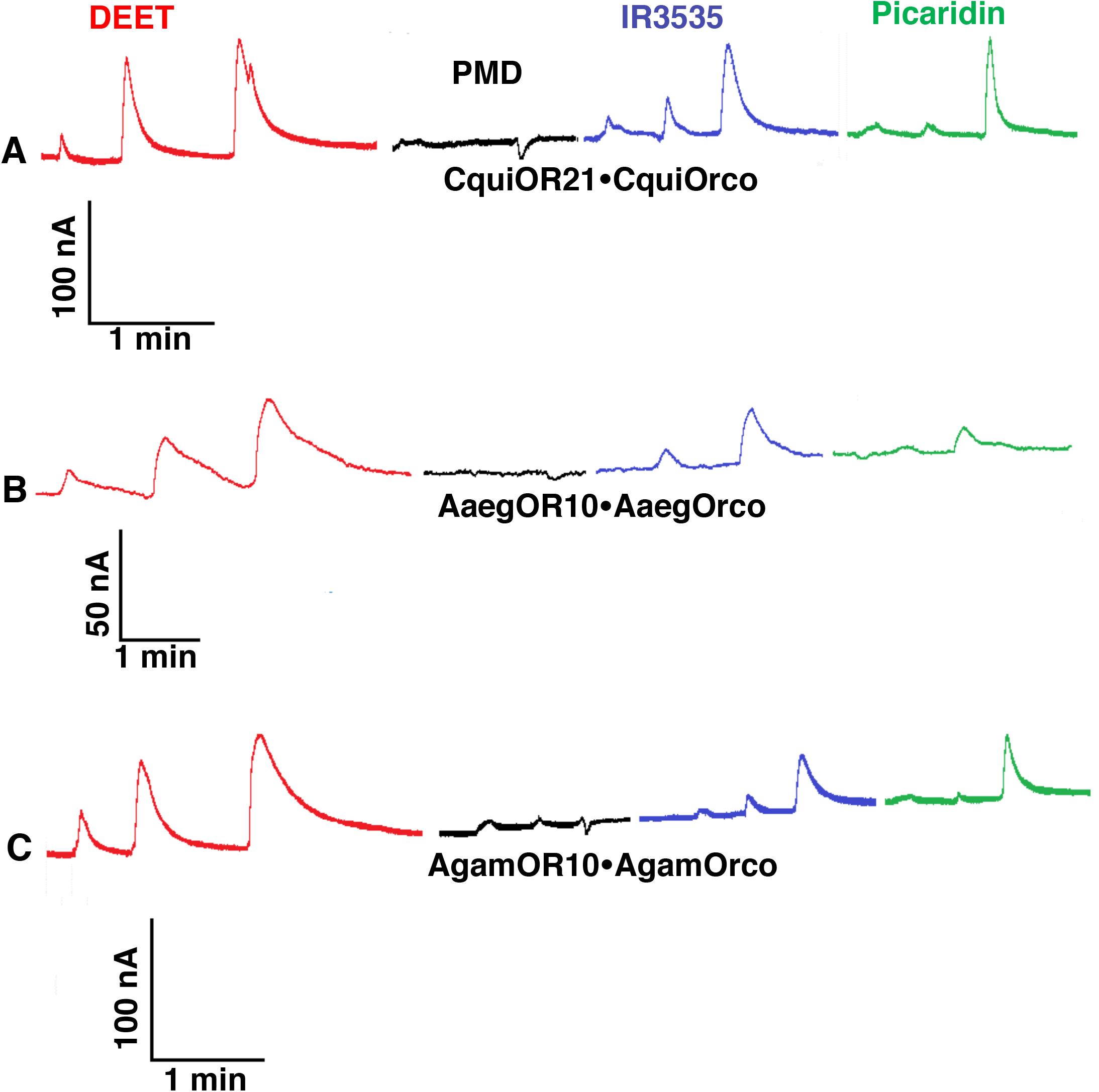
Traces obtained with oocytes expressing CquiOR21 or their orthologues from the yellow fever and malaria mosquito when challenged with repellents. Responses obtained with oocytes co-expressing (A) CquiOR21 and CquiOrco, (B) AaegOR10 and AaegOrco, or (C) AgamOR10 and AgamOrco. All preparations were challenged with DEET (red traces), PMD (black traces), IR3535 (blueberry traces), and picaridin (clover traces) in the same order and with doses 0.01, 0.1, and 1 mM (from left to right).

Next, we challenged CquiOR21, AaegOR10, and AgamOR10 with a panel of 20 compounds, which includes plant-derived and plant-inspired repellents. The compounds are part of pending worldwide (WO2013165477A1) and US (9314029) patent applications and have been previously tested as oviposition deterrents for an agricultural pest, the navel orangeworm, *Amyelois transitella* (Cloonan et al., 2013). The panel was provided to the experimenter (P.X.) with code names, i.e., BDR1-20. To make certain the compounds would be properly identified post hoc, one of us (W.S.L.) analyzed each sample by GC-MS prior to electrophysiology and behavior work.

None of the 20 compounds elicited inward currents (Fig. S4), and 4 compounds did not generate measurable currents, specifically BDR-7, 11, 15, and 20, which were later decoded by W.S.L. to the experimenter. They are ethyl palmitate, 2-pentadecanol, dihydrojasmonic acid, and 2-pentadecanone, respectively. Other compounds generated robust outward currents (equivalent to DEET-elicited currents) at least in one of the three ORs tested. They are BDR-3 (methyl dihydrojasmonate), BDR-4 (methyl jasmonate), BDR-10 (prenyl dihydrojasmonate), BDR-14 (methyl dihydrojasmolate), and BDR-19 (ethyl dihydrojasmonate) (Fig. S4). Of note, repellency activity for methyl jasmonate (Xu et al., 2014) and methyl dihydrojasmonate (Zeng et al., 2018) has been previously demonstrated. Using AgamOR10/AgamOrco-expressing oocytes (Fig. 3), we recorded dose-dependent outward currents generated by these compounds at 0.01, 0.1, and 1 mM.

**Fig. 3.**
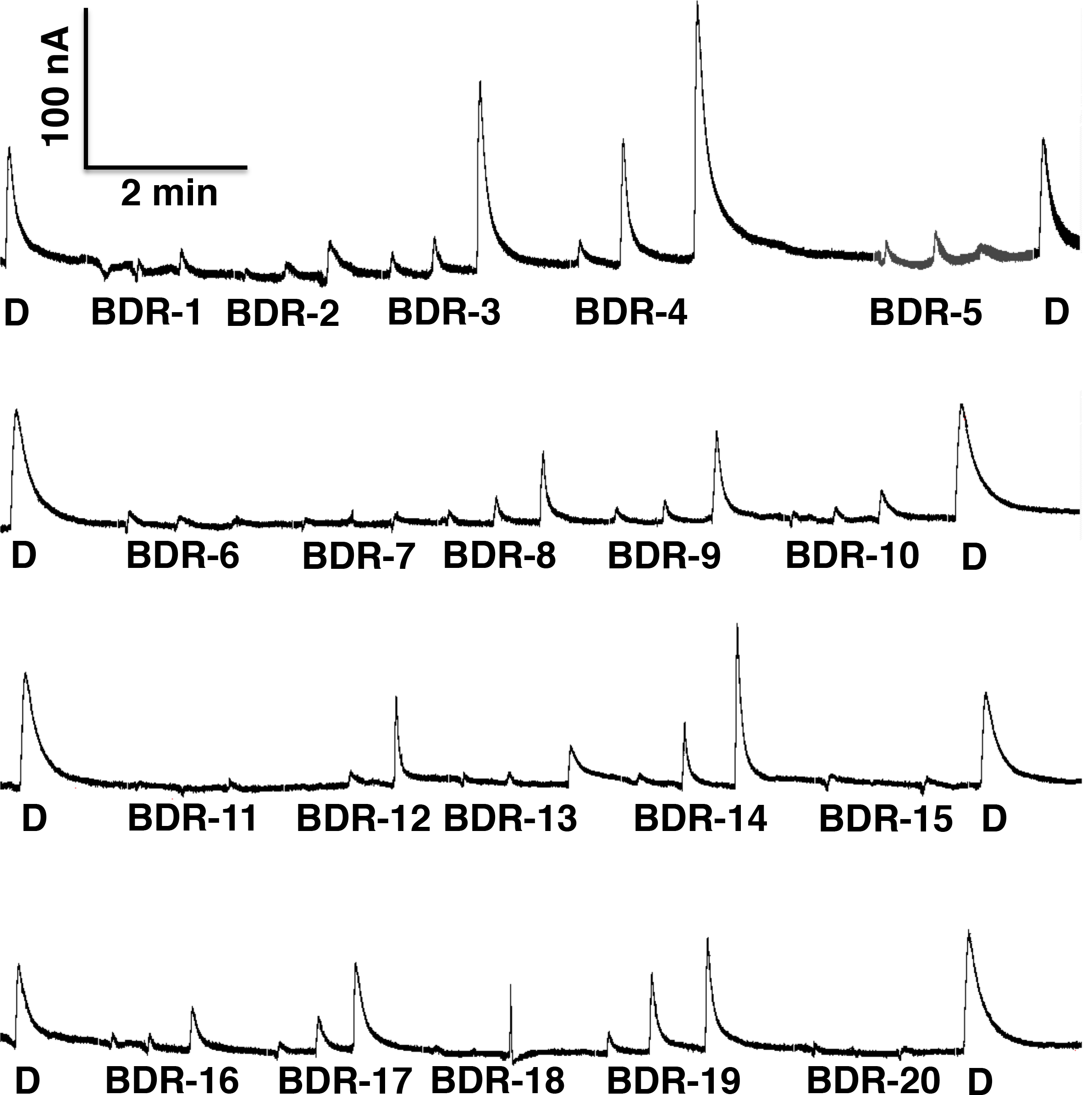
Currents recorded from AgamOR10·AgamOrco-expressing and stimulated with a panel of 20 compounds. DEET (D) was tested at 1 mM and the other compounds (BDR1-20) were tested at 0.01, 0.1, and 1 mM (from left to right). The strongest outward currents were obtained with BDR-4 (methyl jasmonate), BDR-3 (methyl dihydrojasmonate) and BDR-14 (methyl dihydrojasmolate) at 1 mM.

We then investigated whether these outward currents would modulate CquiOR21 responses to skatole. Thus, CquiOR21/CquiOrco-expressing oocytes were challenged with skatole alone or in mixtures with one of the test compounds. Based on preliminary experiments showing that DEET modulates the response to skatole, we selected DEET as a positive control and tested two compounds from our panel, which generated strong/moderate and weak outward currents, i.e., BDR-4 and 5, respectively (Fig. 3, S4). Skatole was presented at a constant dose of 0.1 μM, and the tested compounds were added at decreasing doses from 1 mM to 15 μM (Fig. 4). When mixtures of skatole and DEET or BDR-4 at high doses (1 mM or 0.5 mM) were applied, outward currents were recorded, whereas attenuated inward currents were observed with mixtures containing BDR-5 at the same doses (Fig. 4). The effect of DEET and BDR-4 on CquiOR21 responses to skatole was clearly dose-dependent. When the test compounds were coapplied at 125 μM or lower, only inward currents were recorded. In the case of DEET and BDR-4, the inward currents were attenuated even when these compounds were presented at the lowest dose of 15 μM (Fig. 4). Although this dataset clearly shows that responses to skatole were modulated by DEET (and methyl jasmonate), it does not explain the mode of action of DEET as a noncontact disengagent (= spatial repellent). Mosquitoes responding to CquiOR21 are not host-seeking mosquitoes, but rather gravid females searching for oviposition sites. The observed modulation may explain at least in part the “off-label” activity of DEET as a deterrent for oviposition (Kuthiala et al., 1992). Next, we asked whether compounds modulating OR response to oviposition attractants would activate a DEET receptor mediating spatial repellency (Xu et al., 2014).

**Fig. 4.**
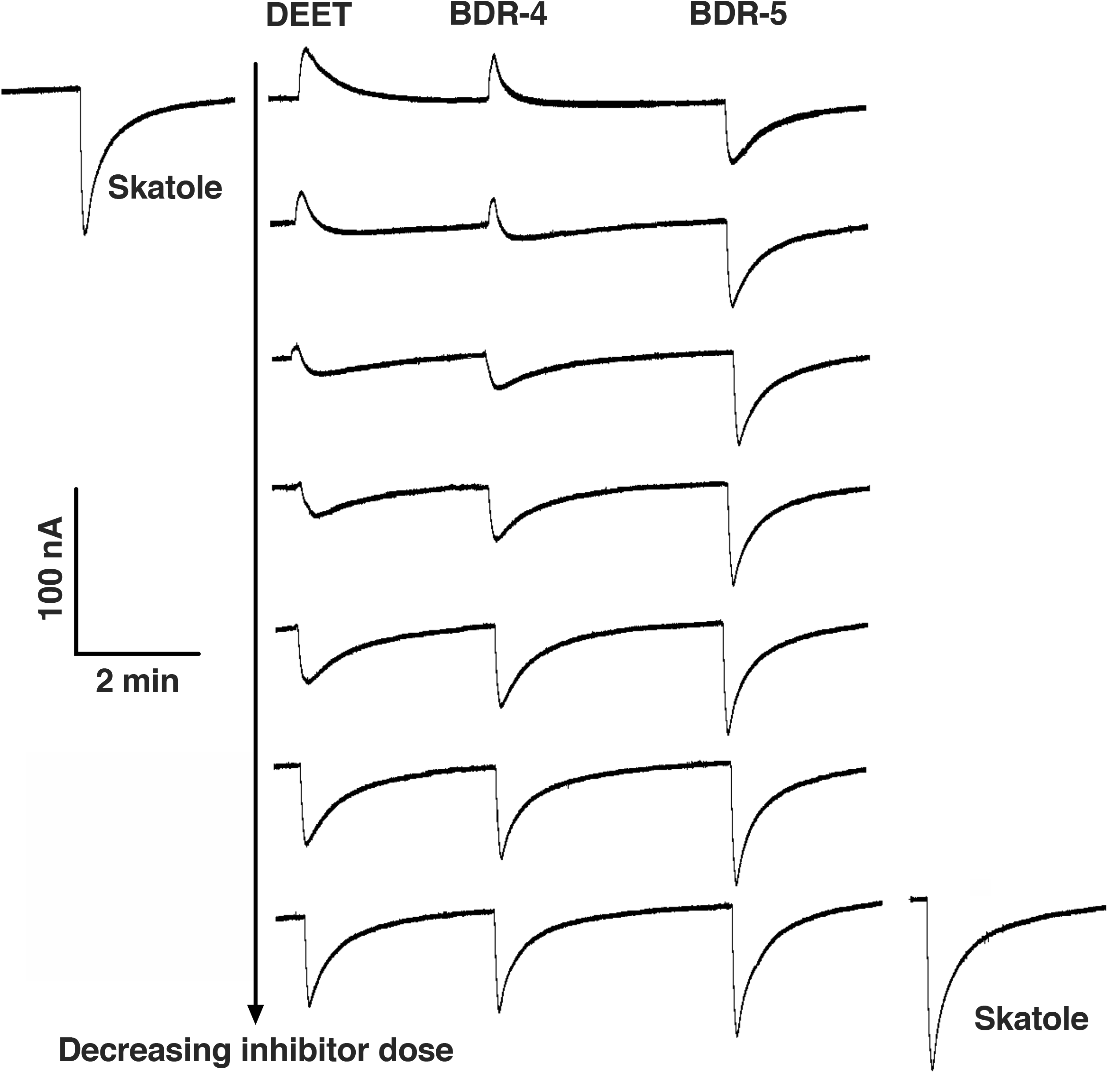
Inhibition of skatole-elicited responses by DEET or methyl dihydrojasmonate. Skatole alone (before: upper left and after: lower right) or in combination with DEET, BDR-4 (methyl jasmonate) or BDR-5 (γ-dodecalactone) was applied at the constant dose of 0.1 μM. DEET, BDR-4, and BDR-5 were applied at 1 mM, 0.5 mM, 0.25 mM, 125 μM, 60 μM, 30 μM, and 15 μM (from top to bottom). At higher doses, the outward currents elicited by DEET or BDR-4 are larger than the inward current generated by skatole, thus generating a net outward current.

### 3.2. Repellent-elicited inward currents

Previously, we have identified CquiOR136 as a DEET receptor in the Southern house mosquito (Xu et al., 2014), which is activated by the four major commercially available repellents, DEET, PMD, IR3535, and picaridin (Xu et al., 2014). CquiOR136/CquiOrco-expressing oocytes were challenged with our panel at three doses (10 μM, 100 μM, and 1 mM) (Fig. S5). IR3535, which elicits the strongest responses at 1 mM (Xu et al., 2014), was used as a positive control. BDR-3 (methyl dihydrojasmonate) and BDR-14 (methyl dihydrojasmolate), among other compounds, elicited robust inward currents (Fig. S5). We then constructed concentration-response relationships for all compounds in our panel (Fig. 5). These analyses clearly show that BDR-14 is the best ligand for CquiOR136 from all tested compounds thus far. More importantly, our data show that CquiOR136 is very sensitive to plant-derived compounds (Fig. 5). Specifically, CquiOR136/CquiOrco-expressing oocytes gave robust responses to methyl dihydrojasmolate, methyl dihydrojasmonate, ethyl dihydrojasmonate, dihydrojasminlactone, dihydrojasmindiol, and methyl jasmonate, which are plant metabolites or their derivatives (plant-inspired compounds). Methyl dihydrojasmolate (BDR-14) is a reduced form of methyl dihydrojasmonate (hydroxy vs. a carbonyl moiety), which in turn is the product of hydrogenation of the plant hormone methyl jasmonate. That this DEET receptor is very sensitive to these plant-derived and plant-inspired compounds is consistent with the notion that the primary function of CquiOR136 in the biology of *Cx. quinquefasciatus* is the reception of plant defense compounds and that DEET mimics these natural products (Xu et al., 2014).

**Fig. 5.**
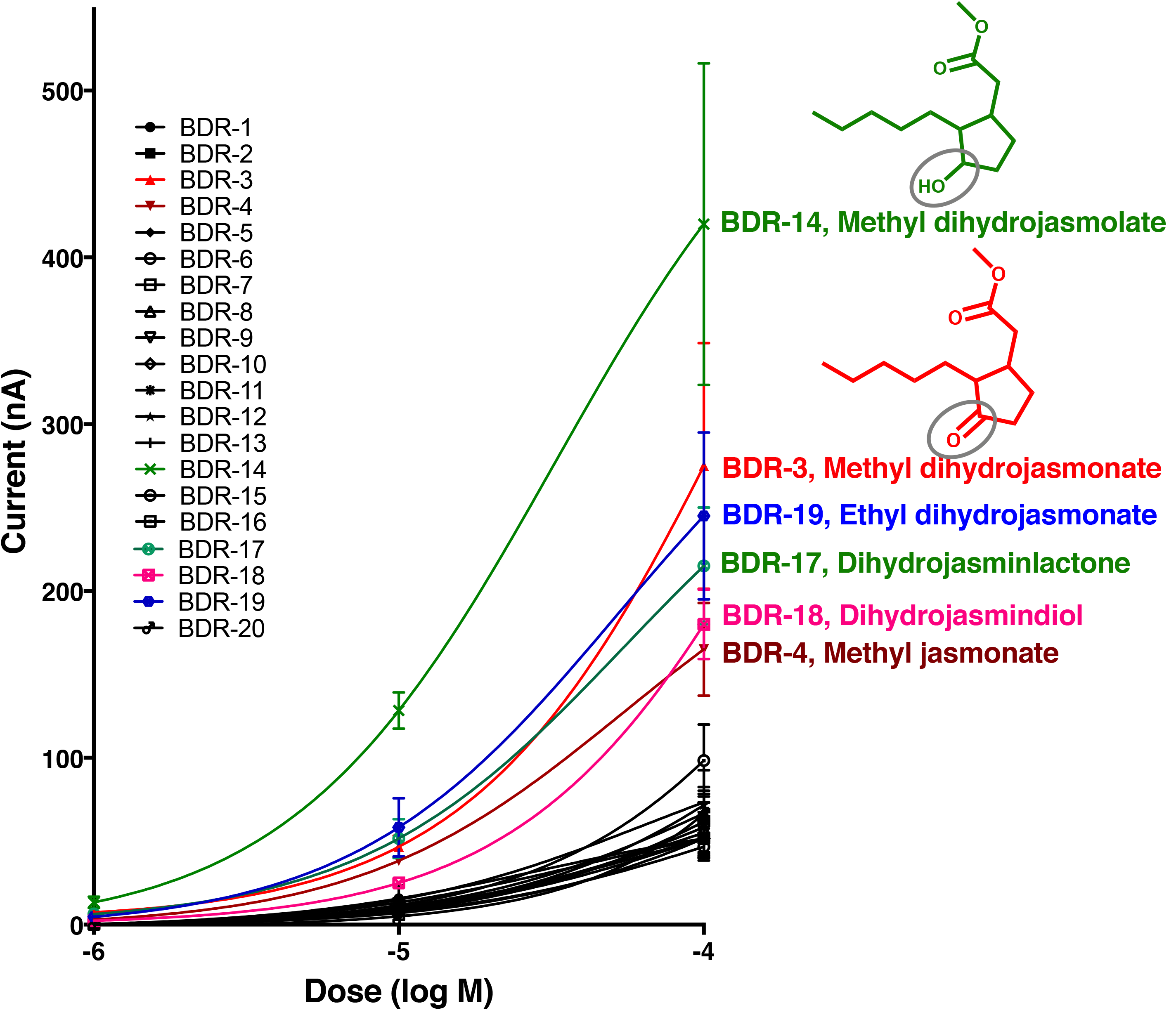
Concentration-response relationships of CquiOR136·CquiOrco in response to test repellents. Mean ± SEM, *N* = 3 for each point. Data obtained with different oocytes were not normalized.

### 3.3. Repellency activity of methyl dihydrojasmolate

Given the robust responses recorded of CquiOR136 to methyl dihydrojasmolate, we tested the repellency activity of this compound using our surface landing and feeding assay (Leal et al., 2017; Xu et al., 2014). First, we compared the repellency activity of methyl dihydrojasmolate to DEET with both compounds at 0.1%. At this dose, DEET showed ca. 80% protection, whereas no protection was achieved with methyl dihydrojasmolate (n = 5 each, unpaired, two-tailed *t* test, P = 0.0020) (Fig. 6A). At 1% dose, methyl dihydrojasmolate gave almost 60% protection, but significantly lower activity than DEET (n = 5 each, Mann-Whitney two-tailed test, P=0.0088) (Fig. 6B). We surmised that the lower protection rate obtained with methyl dihydrojasmolate might be due to differences in volatility. Measurements of spatial repellency are biased by differences in vapor pressures, with compounds with lower vapor pressure yielding lower protection, but longer duration. DEET has a higher vapor pressure than methyl dihydrojasmolate. Therefore, we compared methyl dihydrojasmolate at a higher dose (10%) with 1% DEET. Even with our attempt to compensate for vapor pressure, DEET showed a significantly better performance (n = 5 each, unpaired, two-tailed *t* test, P = 0.0445) (Fig. 6C). These findings suggest that comparatively DEET is a better spatial repellent, but we cannot unambiguously conclude whether DEET would have a better overall performance because high contact repellency activity may compensate for moderate spatial repellency.

**Fig. 6.**
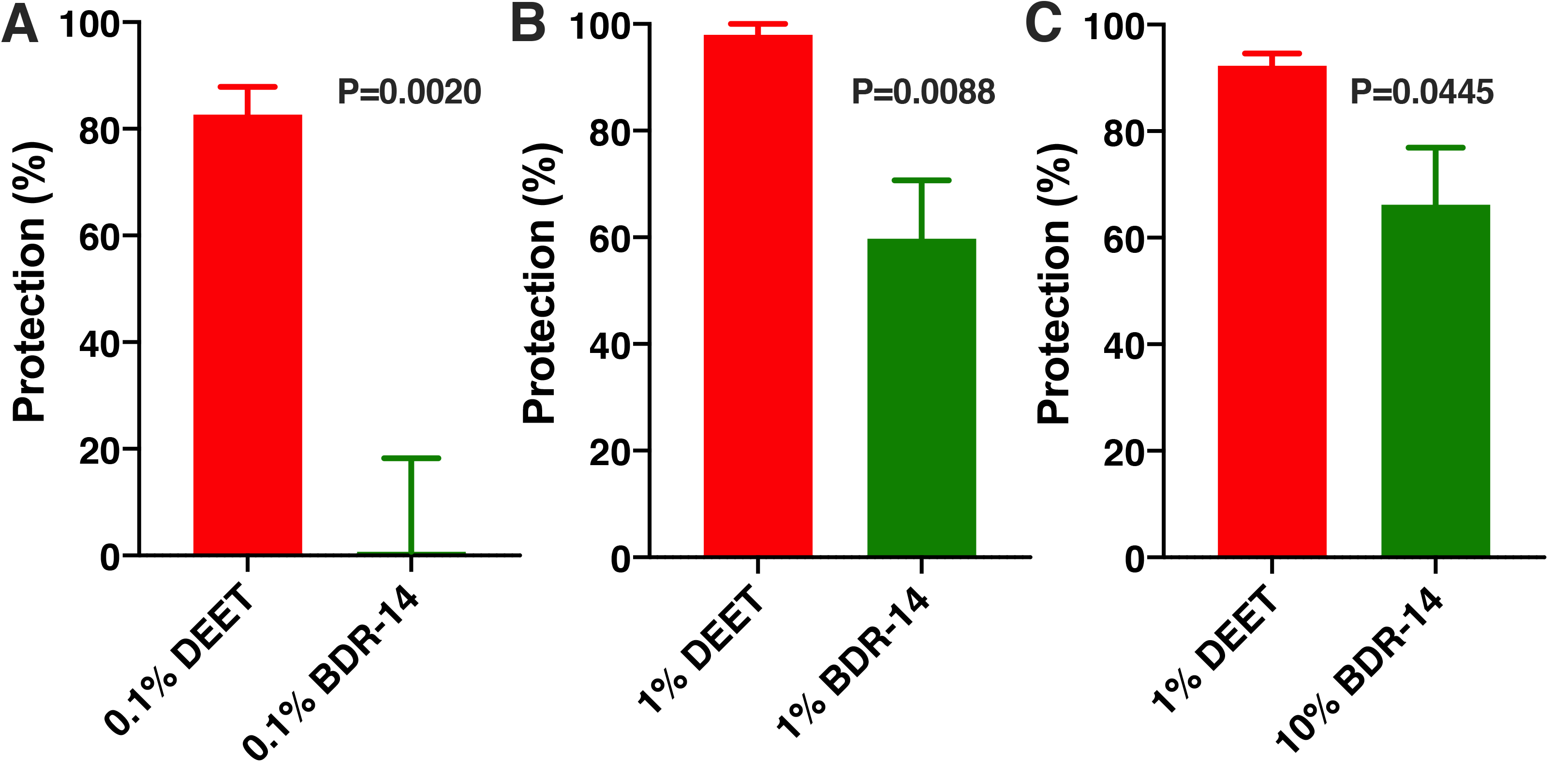
Repellency activity of DEET and methyl dihydrojasmolate. Repellency was tested against the Southern house mosquito and expressed in protection rate using the following doses: (A) both compounds at 0.1%, (B) both compounds at 1%, and (C) 1% DEET and 10% methyl dihydrojasmolate. n = 5 for each test compound at each dose.

### 3.4. Overall conclusions

Our data suggest that mosquito response to oviposition attractants may be modulated by repellents. When ORs sensitive to oviposition attractants were challenged with repellent, outward (inhibitory, hyperpolarizing) currents were generated. Responses of the OR detecting the oviposition skatole in the Southern house mosquito, CquiOR21 (=CquiOR10), were reduced when skatole was coapplied with DEET or methyl dihydrojasmolate. These inhibitory currents may explain at least in part the deterrent effect of DEET on attraction of gravid females (Kuthiala et al., 1992). More importantly, it demonstrates that integration of chemical signals at the peripheral olfactory system is more complex than previously appreciated.

## Acknowledgments

The authors are grateful to Dr. Laurence (Larry) J. Zwiebel, Vanderbilt University, for sharing AgamOrco, AgamOR10, AgamOR8, AaegOR10 plasmids, and Dr. Anthon J. Cornel, UC Davis, Department of Entomology and Nematology, for providing *Ae. aegypti* mosquitoes for RNA extraction.

## Author Contributions

P.X. and W.S.L. conceived the project, designed the experiments, and performed data analysis. P.X. performed electrophysiology. F.Z. measured mosquito behavior. W.S.L. performed GC-MS analysis. R.H.B. prepared and provided test compounds. W.S.L. wrote the manuscript. All authors provided input, read, and approved the final version of the manuscript.

## Supplementary information

**Fig. S1.**
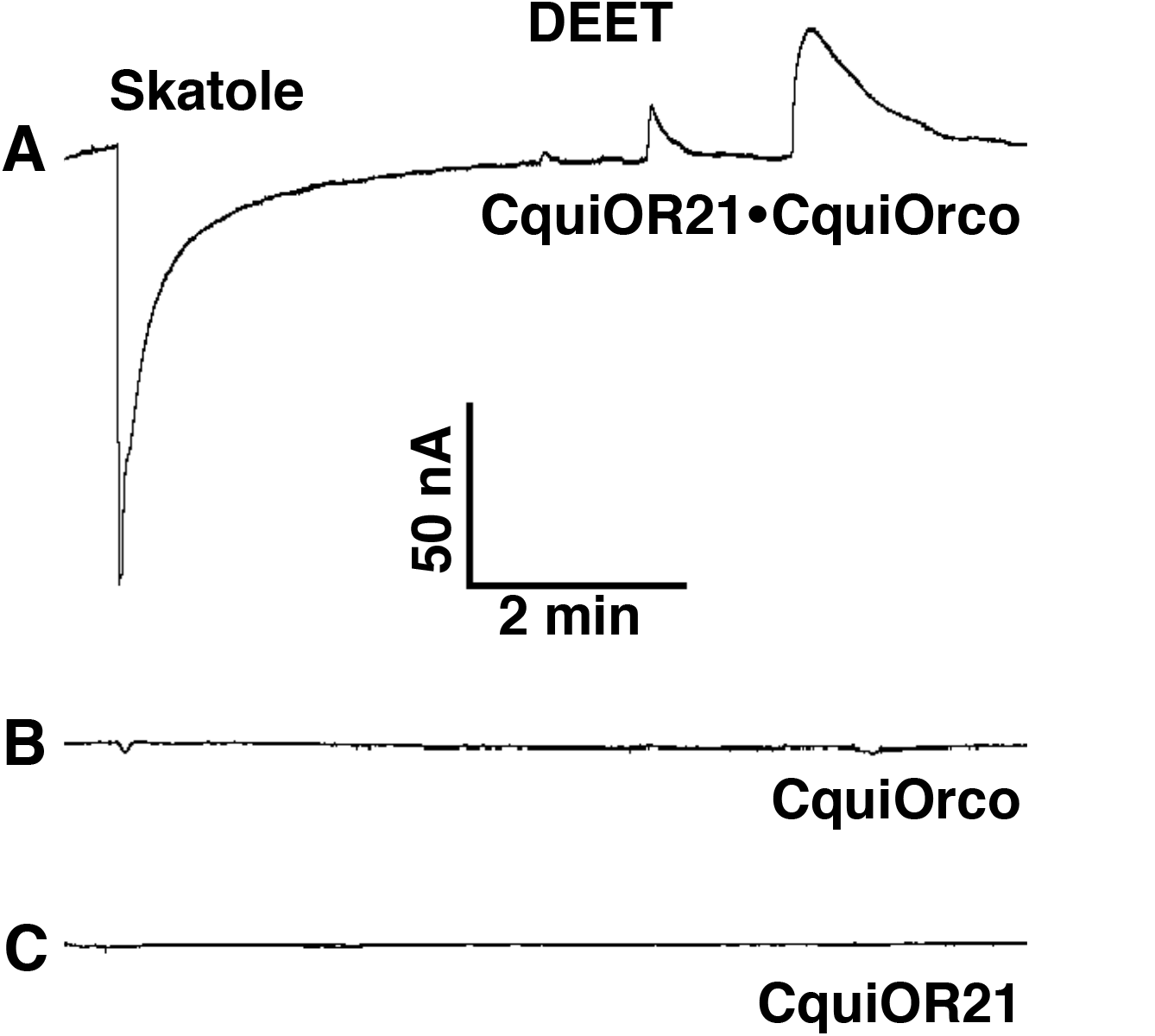
Responses of a CquiOR21·CquiOrco-expressing oocyte to DEET and skatole. **(A)** Inward current elicited by 1 μM skatole and outward currents generated by DEET at 0.01, 0.1, and 1 mM (from left to right). Test of control oocytes injected with (B) CquiOrco or (C) CquiOR21 challenged with the same compounds at the same doses.

**Fig. S2.**
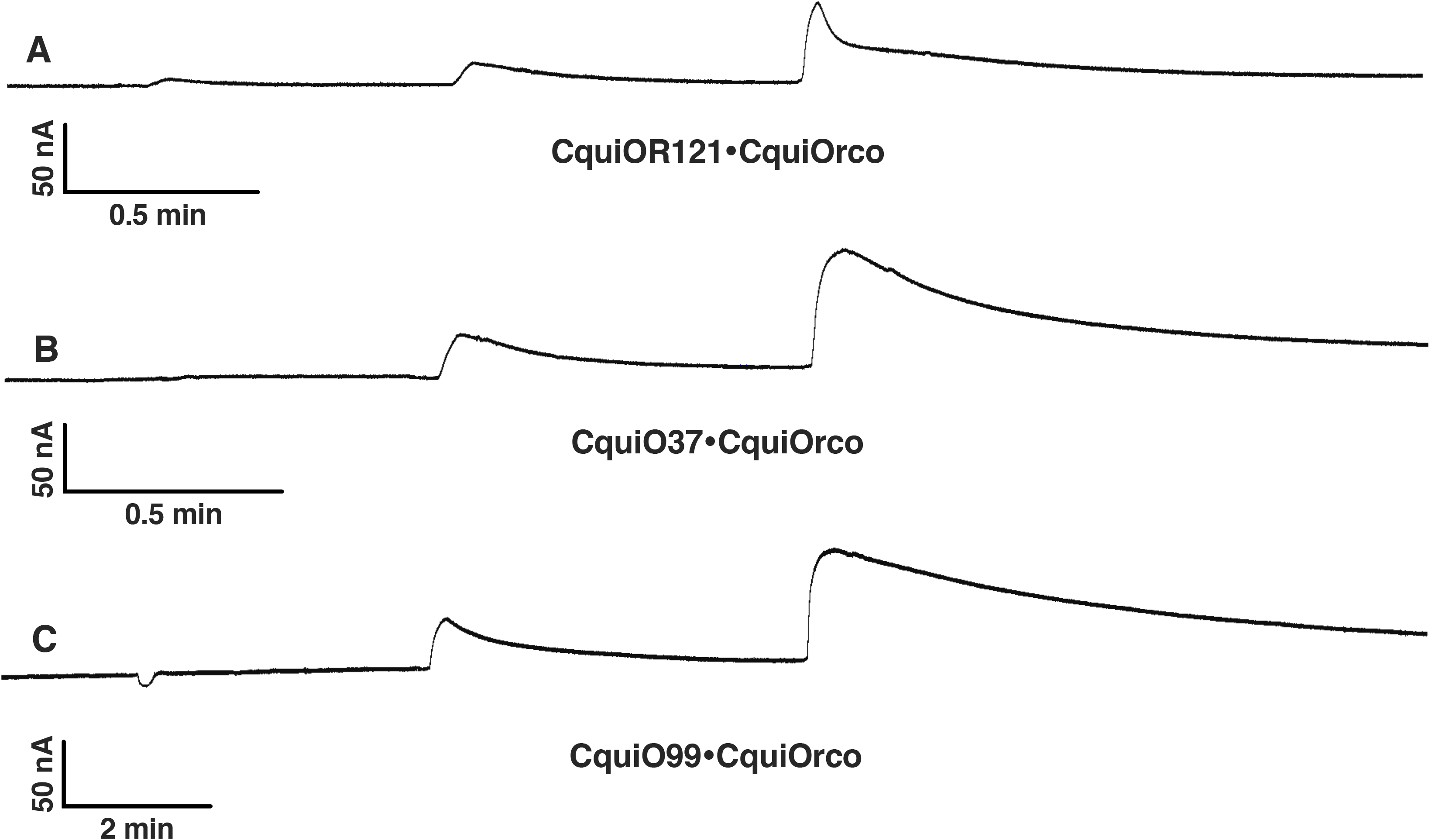
Outward currents elicited by DEET on other oviposition attractant ORs. DEET was applied at 0.01, 0.1, and 1 mM (from left to right). Responses elicited by (A) CquiOR121, (B) CquiOR37, and (C) CquiOR99.

**Fig. S3.**
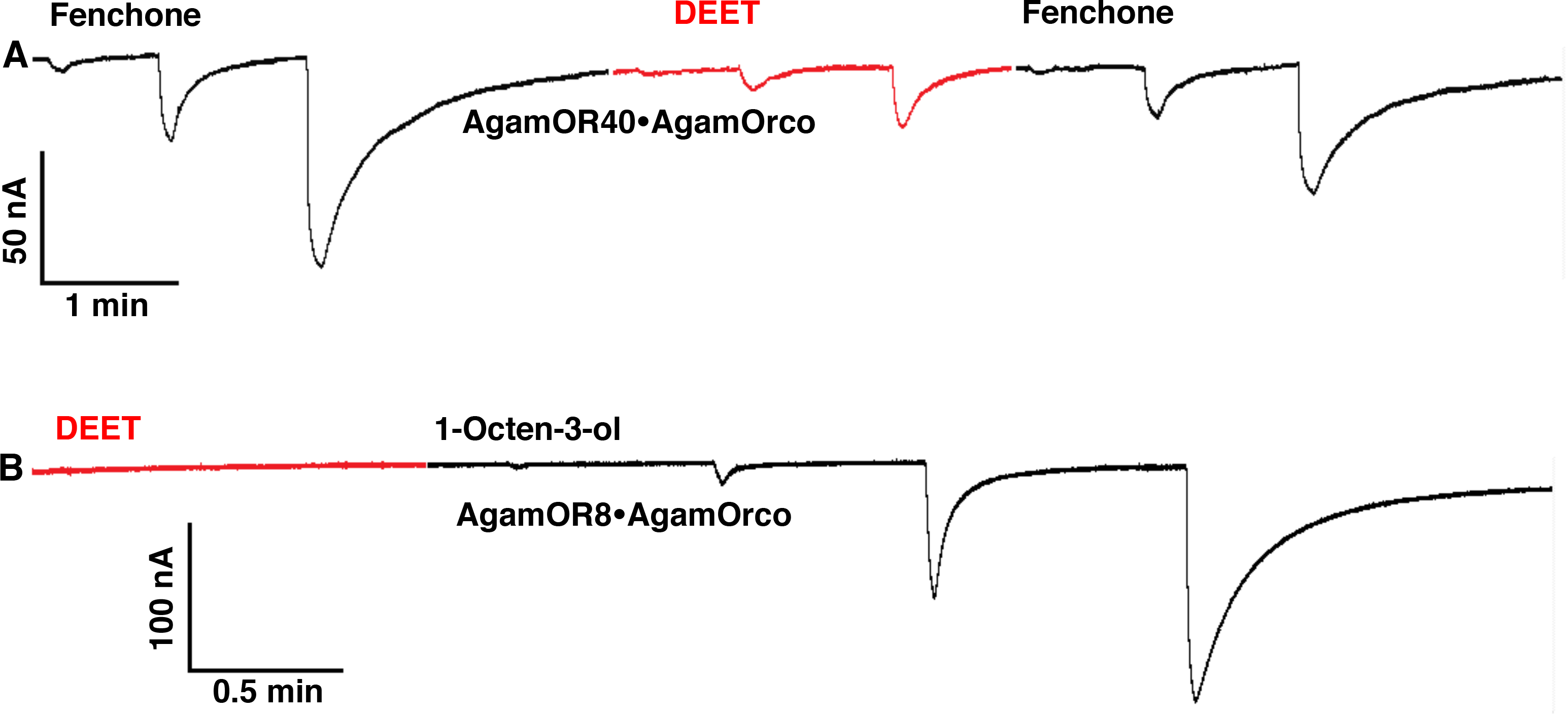
Inward currents and no response recorded from other ORs challenged with DEET. (A) Responses recorded from an oocyte expressing the larval odorant receptor, AgamOR40, along with its coreceptor AgamOrco. Fenchone (black traces) and DEET (red trace), 0.01, 0.1., and 1 mM (from left to right). (B) Currents recorded from an oocyte coexpressing AgamOR8 and AgamOrco and challenged with DEET (red trace, 0.01, 0.1, and 1 mM) and 1-octen-3-ol (black trace: 0.1 and 1 μM, 0.01 and 0.1 mM from left to right).

**Fig. S4.**
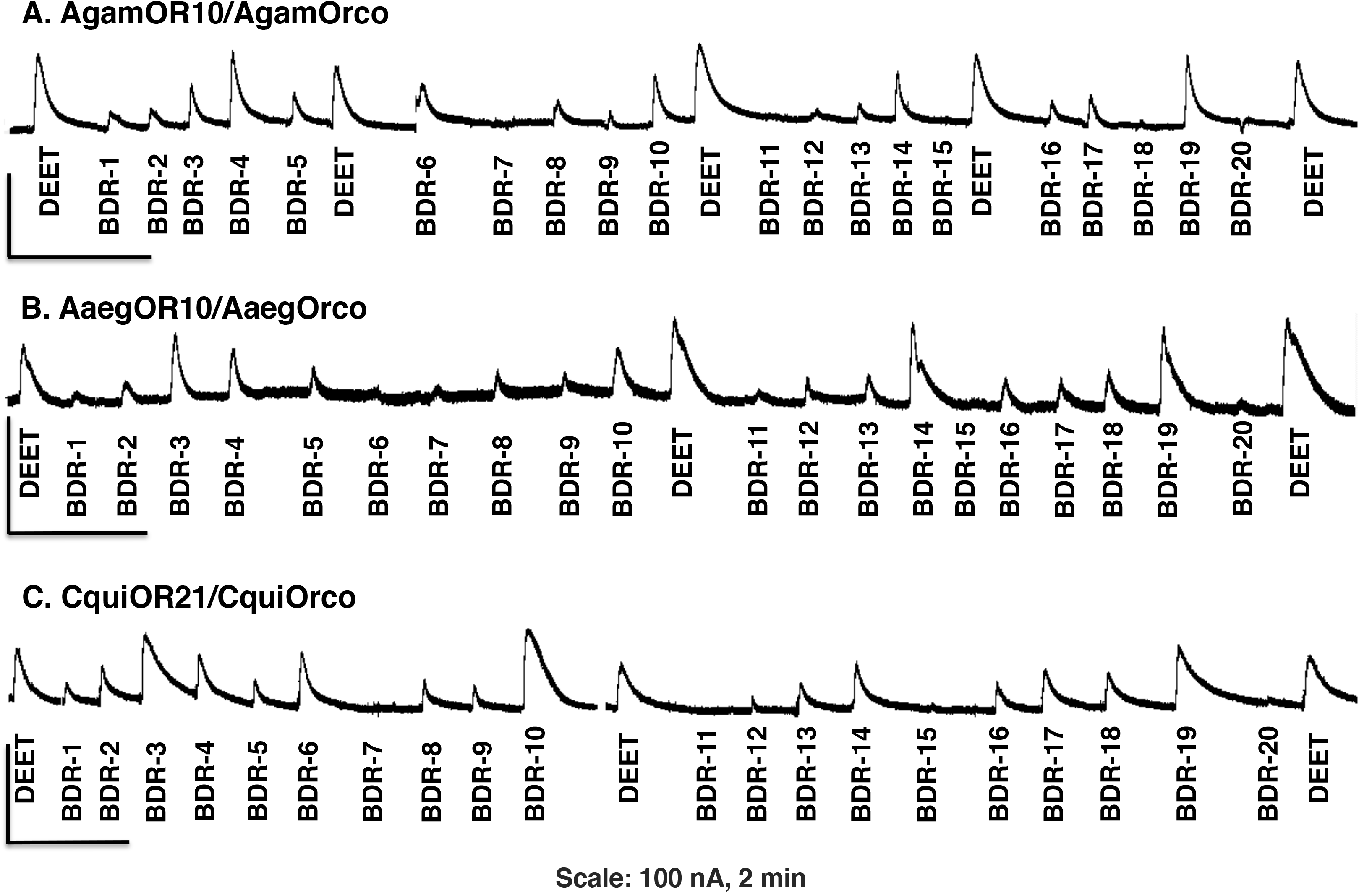
Comparison of outward currents recorded from OR orthologues. Responses elicited by compounds BDR-1 to BDR-20 on oocytes expressing (A) AgamOR10 and AgamOrco, (B) AaegOR10 and AaegOrco, and (C) CquiOR21 and CquiOrco. All compounds, including DEET, were tested at 1 mM. The profiles obtained with the three species are slightly different. In general, BDR-1, 2, 5, 7, 8, 9, 11, 12, 13, 15, 16, 17, 18, and 20 showed weak or no response. By contrast, BDR-3, 4, 10 (prenyl dihydrojasmonate), 14, and 19 (ethyl dihydrojasmonate) elicited moderate or robust outward currents.

**Fig. S5.**
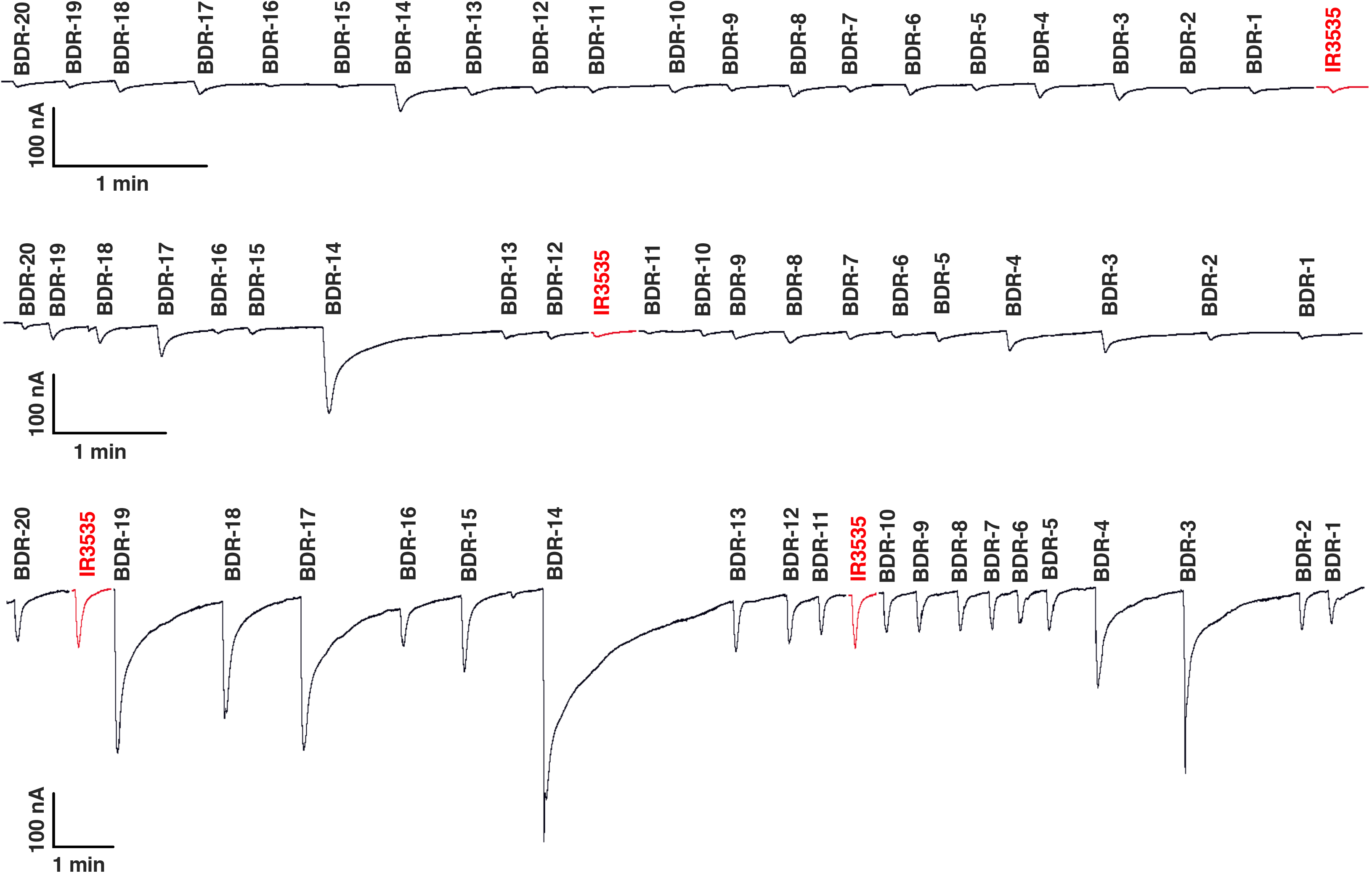
Traces obtained with CquiOR136/CquiOrco-expressing oocytes. Compounds were presented at 1 μM (upper trace), 10 μM (middle trace), and 100 μM (bottom trace).

